# GABA_B_ Receptors Gate Sex-Specific Synaptic Plasticity in the Nucleus Accumbens

**DOI:** 10.64898/2026.08.26.747391

**Authors:** Ashley E. Copenhaver, Tara A. LeGates

## Abstract

Excitatory synaptic plasticity within the nucleus accumbens (NAc) drives motivated behaviors, and dysregulation is implicated in several psychiatric disorders marked by impaired reward processing. The NAc integrates glutamatergic input, which conveys information about reward, context, and behavioral goals, with local GABAergic signaling that regulates excitatory transmission and medium spiny neuron (MSNs) output. However, little is known regarding GABA-dependent modulation of activity-dependent excitatory synaptic plasticity. Here, we investigated GABA_B_ receptor (GABA_B_R) regulation of plasticity at hippocampus (Hipp)-NAc synapses, at which plasticity is a key mediator of reward-related behaviors. Using whole-cell electrophysiological recordings in mouse brain slices, we found that pharmacological inhibition of GABA_B_Rs converts long-term potentiation (LTP) into long-term depression (LTD) selectively in females, identifying a sex-specific role for GABA_B_Rs in modulating long-term plasticity of Hipp-MSN synapses. This LTD required mGluR5 activation and estrogen receptor alpha (ERα) in both D1- and D2-expressing MSN subtypes, while only D1-MSNs suggested that LTD was expressed presynaptically through a CB1 receptor-dependent mechanism. Notably, GABA_B_R inhibition did not alter basal synaptic transmission, indicating a specific role for these receptors in gating plasticity beyond regulation of basal excitatory drive. Together, these findings identify a novel, sex-specific mechanism by which GABA_B_Rs control the direction of synaptic plasticity.

## 1. Introduction

Excitatory synaptic plasticity in the reward system supports experience-dependent changes in reward processing and motivated behavior, and alterations are associated with several psychiatric disorders characterized by reward-related dysfunction, including depression and substance use disorders (SUDs)[1]. The nucleus accumbens (NAc), a central node of the reward system, integrates excitatory inputs from several upstream brain areas that convey key information relevant to goal-directed behaviors. Modulation of excitatory synaptic strength has a major influence on how the NAc translates reward-related experiences into adaptive behavioral outputs[2-3].

Within the NAc, the integration of local GABAergic inhibitory networks is critical for modulating excitatory transmission and influence circuit activity and behavior. GABA_B_Rs are well-positioned to modulate excitatory synaptic transmission through both pre- and postsynaptic mechanisms and have been implicated in reward processing and anhedonia[4-6]. Recent work has shown that GABA_B_Rs gate excitatory transmission within the NAc[6], underscoring their potential to regulate reward circuit plasticity in a pathway-specific manner. However, there is little understanding of how GABAergic modulation, particularly through GABA_B_ receptors (GABA_B_Rs), impacts excitatory synaptic plasticity.

Here, we tested whether GABA_B_Rs influence long-lasting synaptic plasticity at hippocampus (Hipp)-NAc synapses. Excitatory input from the Hipp drives NAc neuron activity, and Hipp-NAc synapses are a key site where environmental context is linked with reward-related outcomes to support the proper execution of goal-directed behaviors[7-12]. The bidirectional modulation of Hipp-NAc synaptic strength is a critical mediator of motivated behaviors, where potentiation is rewarding and necessary for reward learning while weakening induces anhedonia and learning deficits[13]. We show that pharmacological inhibition of GABA_B_Rs switches the sign of plasticity in female but not male mice, revealing a previously unrecognized sex-specific role for GABA_B_Rs in controlling the direction of excitatory synaptic plasticity.

## 2. Materials and Methods

### 2.1 Animals

8-10-week-old D1dra-tdTomato or C57BL/6J mice were bred in-house. Mice were housed with same-sex cage mates in a temperature- and humidity-controlled environment under a 12:12h light/dark cycle (lights on at 07:00). We did not track the estrous cycle. All experiments were performed in accordance with the regulations set forth by the Institutional Animal Care and Use Committee at the University of Maryland, Baltimore County.

### 2.2 Mouse brain slice preparation for electrophysiology

Acute parasagittal slices (lateral 0.36-0.72) containing the fornix and nucleus accumbens were prepared for whole-cell patch-clamp electrophysiology. Animals were deeply anesthetized with isoflurane, and brains were quickly dissected and submerged in ice-cold, oxygenated *N*-methyl-D-glucamine (NMDG) recovery solution containing (in mM): 93 NMDG, 2.5 KCl, 1.2 NaH_2_PO_4_, 11 glucose, 25 NaHCO_3_, 1.2 MgCl_2_, and 2.4 CaCl_2_). Parasagittal slices (400μm) were cut on a vibratome (VT1000S, Leica Microsystems) in cold, oxygenated NMDG. Slices were transferred to 32–34°C NMDG for 7-12 min and then transferred to room temperature artificial cerebrospinal fluid (aCSF) containing (in mM): 120 NaCl, 3 KCl, 1.0 NaH2PO4, 20 glucose, 25 NaHCO3, 1.5 MgCl2·7H2O, and 2.5 CaCl2, pH 7.3–7.4. Slices recovered for 1 hour at room temperature before electrophysiological recordings.

### 2.3 Whole-cell recordings

We performed whole-cell patch-clamp recordings using an Axopatch 200B amplifier and Digidata 1550B digitizer (Molecular Devices). Slices were placed in a submersion-type recording chamber and superfused with room temperature aCSF (flow rate, 0.5-1 ml/min) and visualized using a 60× water immersion objective (Nikon Eclipse FN-1). Patch pipettes (4-8 MΩ) were made from borosilicate glass (World Precision Instruments) using a Sutter Instruments P-97 model puller. Dopamine-1-receptor and putative dopamine-2-receptor-expressing MSNs (D1-MSN and pD2-MSN) were identified by the presence and absence of tdTomato expression, respectively.

A bipolar stimulating electrode (FHC) was placed in the fornix to electrically stimulate hippocampal axons, and excitatory postsynaptic currents (EPSC) were recorded in the NAc medial shell. Patch pipettes were filled with a solution containing 130mM K-gluconate, 5mM KCl, 2mM MgCl_6_-H_2_O, 10mM HEPES, 4mM Mg-ATP, 0.3mM Na_2_-GTP, 10mM Na_2_-phosphocreatine, and 1mM EGTA; pH=7.3-7.4; 285-295mOsm. For LTP, a 5-minute baseline EPSC recording was obtained from paired pulses (100ms ISI) elicited every 10s. This was followed by high-frequency stimulation (HFS: four trains of 100Hz stimulation for 1s with 15s between trains at -40mV), and paired-pulses were recorded again for 30-minutes. For pharmacological manipulation, ACSF-containing drugs (or vehicle for controls) were superfused for at least 15 minutes prior to recording at the following concentrations: SCH50911 (Tocris, 5μM), AM251 (Tocris, 3μM), MPEP hydrochloride (Tocris, 10μM), MPP dihydrochloride (Tocris, 3μM). For wash-on experiments, a stable baseline was recorded followed by wash-on of SCH50911 or vehicle-containing ACSF while paired pulse stimulation continued. Cells were excluded if: 1) series resistance >10MΩ, 2) series resistance changed by >20% (comparing the resistance at the beginning and end of the experiment), 3) Poor health or poor recording status (i.e. cell partially or fully sealed up, a decrease in holding current >100pA that is consistent with the cell dying, and/or an increase in jitter post HFS).

### 2.4 Data analysis

EPSC amplitudes were normalized to the mean baseline amplitudes for comparison across cells. EPSCs are plotted in one-minute bins (mean +/-SEM). Mean EPSC amplitudes during the last 5 minutes of recording were used for statistical analysis. Paired-pulse ratio (PPR) was determined by calculating 5-min averages of EPSC amplitudes from our paired pulses (EPSC1 and EPSC2) and dividing EPSC2 by EPSC1. Change in PPR was calculated as the difference between the PPR at 25–30 min post-HFS and baseline PPR, normalized to baseline PPR. For box plots, the line in the middle of the box is plotted at the median. The box extends from the 25th to 75th percentiles. Whiskers represent minimum and maximum, and individual points are shown. All statistical analyses were performed using GraphPad Prism 10.3.0. Statistics for group comparisons are described in the corresponding figure legends. One-sample t-test or Wilcoxon tests were used to determine whether EPSC amplitudes were significantly different from baseline.

## 3. Results

### 3.1 Inhibition of GABABRs results in sex-specific sign-change in long-lasting plasticity

We pretreated slices with the GABA_B_R antagonist SCH50911 and found that HFS induced LTD in females (Figure 1a-b), a marked difference from both control conditions and our prior findings in which HFS reliably induced LTP (Figure 1a-b)[13-14]. Analysis of paired-pulse ratio (PPR) revealed an increase in PPR selectively in D1-MSNs, suggesting a potential presynaptic contribution to LTD in this cell-type (Figure 1a,c). Interestingly, these effects on plasticity were not observed in males (Figure 1d-f). Together, these findings identify a novel, sex-specific role for GABA_B_Rs in modulating the direction of excitatory synaptic plasticity.

**Figure 1:**
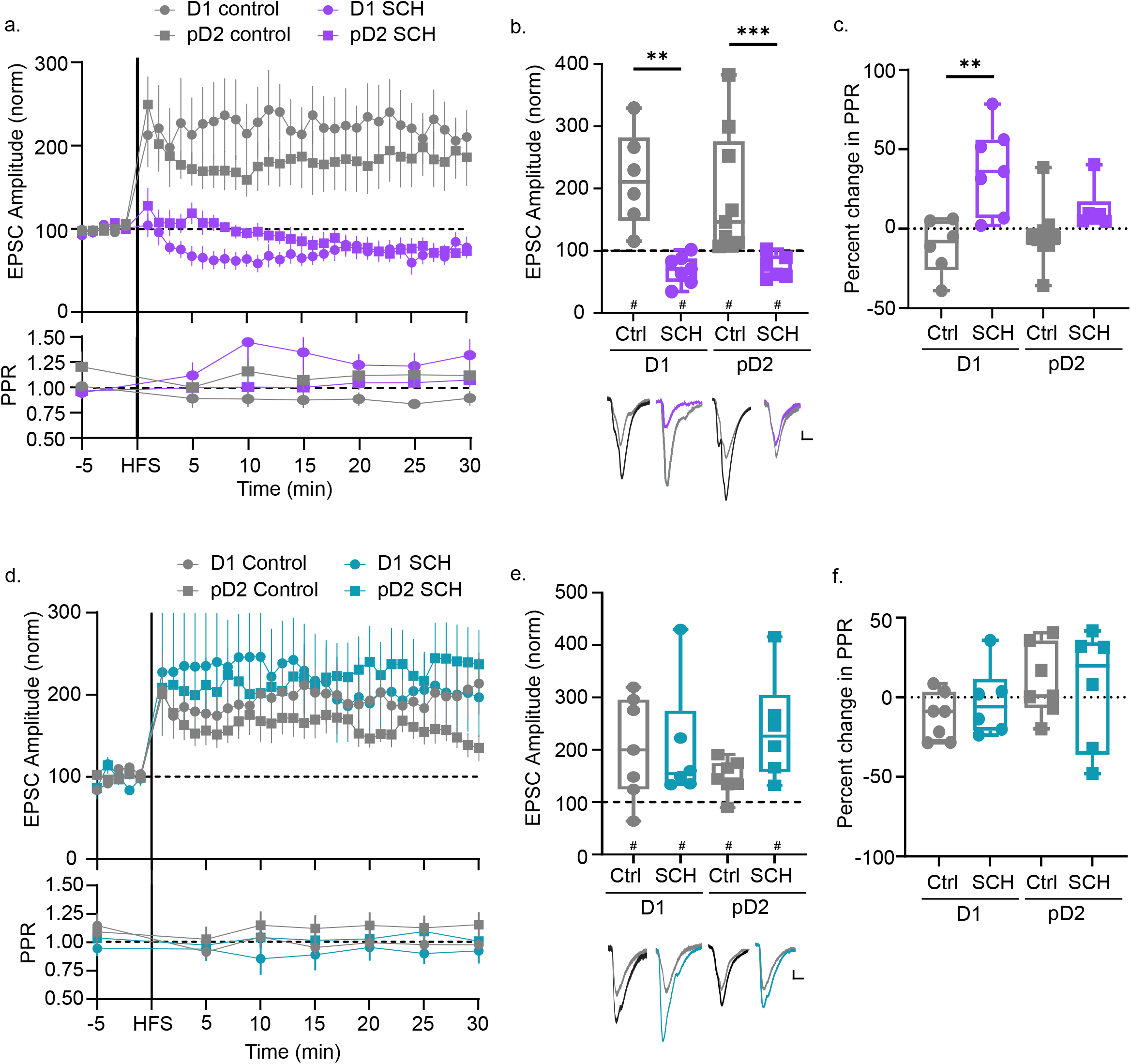
GABA_B_Rs inhibition changes the sign of HFS-induced plasticity in females. a) EPSCs and PPR before and after HFS in the presence and absence of SCH50911 (SCH) in D1- and pD2-MSNs from females. b) Average EPSC amplitude post-HFS with representative traces. c) Comparison of the percent change in PPR during the HFS experiment. (Control D1 F n=6 cells/6 mice, W=21, ^#^p=0.0312; SCH D1 F n=7 cells/7 mice, W=-26, ^#^p=0.0312; Control pD2 F n=9 cells/9 mice, W=45, ^#^p=0.0039; SCH pD2 F=7 cells/6 mice, W=-26, ^#^p=0.0312) d) EPSCs and PPR before and after HFS in the presence and absence of SCH50911 in D1- and pD2-MSNs from males. e) Average EPSC amplitude post-HFS with representative traces. f) Comparison of the percent change in PPR. (Control D1 M n=7 cells/7 mice, W=24, p=^#^0.0469; SCH D1 M n=6 cells/6 mice, W=21, p=^#^0.0312; Control pD2 M n=7 cells/7 mice, W=26, p=^#^0.0312; SCH pD2 M=6 cells/6 mice, W=21, p=^#^0.0312). EPSC amplitude: Three-way ANOVA:F_(1,47)(sex x treatment)_=15.36; F_(1,47)(treatment)_=4.900; F_(1,47)(sex)_=9.794; Šídák’s:Female D1_control vs SCH_ *p=0.0238, Female D2_control vs SCH_ *p=0.0498. PPR: Three-way ANOVA:F_(1,40)(sex x treatment)_=5.490; F_(1,40)(cell-type x sex)_=4.462; F_(1,40)(cell-type x treatment)_=4.083; F_(1,40)(treatment)_=7.686; Šídák’s:*p_(Female D1 control vs SCH)_=0.0033, *p_(Female D1 SCH vs Male D1 SCH)_=0.0241. Scalebars: 20pA/10ms.

To determine whether the LTD that occurs during GABA_B_R blockade (herein referred to as SCH-LTD) was due to GABA_B_ receptor-dependent modulation of basal transmission, we recorded EPSCs in slices taken from females during wash-on of SCH50911. We found that SCH50911, similar to vehicle, did not alter EPSC amplitude or PPR (Figure 2a-b), suggesting that SCH-LTD is not attributable to changes in basal synaptic transmission or presynaptic release probability.

**Figure 2:**
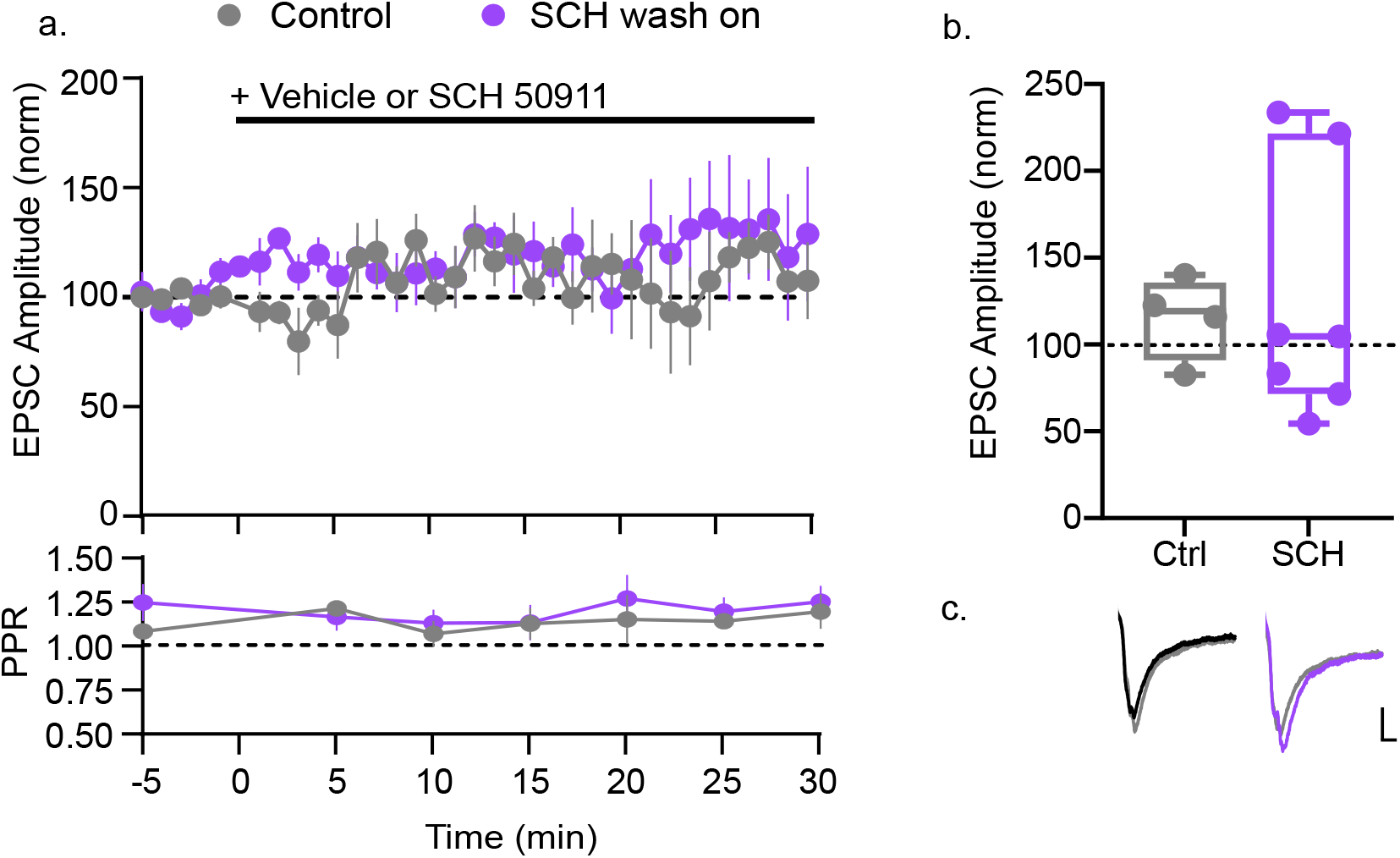
GABA_B_Rs inhibition does not alter basal synaptic function. a) EPSC amplitude and PPR during wash-on of SCH50911 in females. b) Average EPSC amplitude 25-30min post-wash-on with representative traces. (Control n=4 cells/4 mice, SCH n=7 cells/4 mice. Mann Whitney U: U=11, p=0.6485). Scalebars: 20pA/10ms.

### 3.2 SCH-LTD Mechanism Interrogation

To determine the mechanisms underlying SCH-LTD that was observed in females, we first tested the involvement of endocannabinoid signaling, a prominent mediator of LTD[15]. We found that the CB1 receptor (CB1R) antagonist, AM251, prevented SCH-LTD selectively in D1-MSNs (Figure 3a-b), revealing a cell type-specific requirement for CB1R signaling consistent with our PPR data (Figure 2a-b).

**Figure 3:**
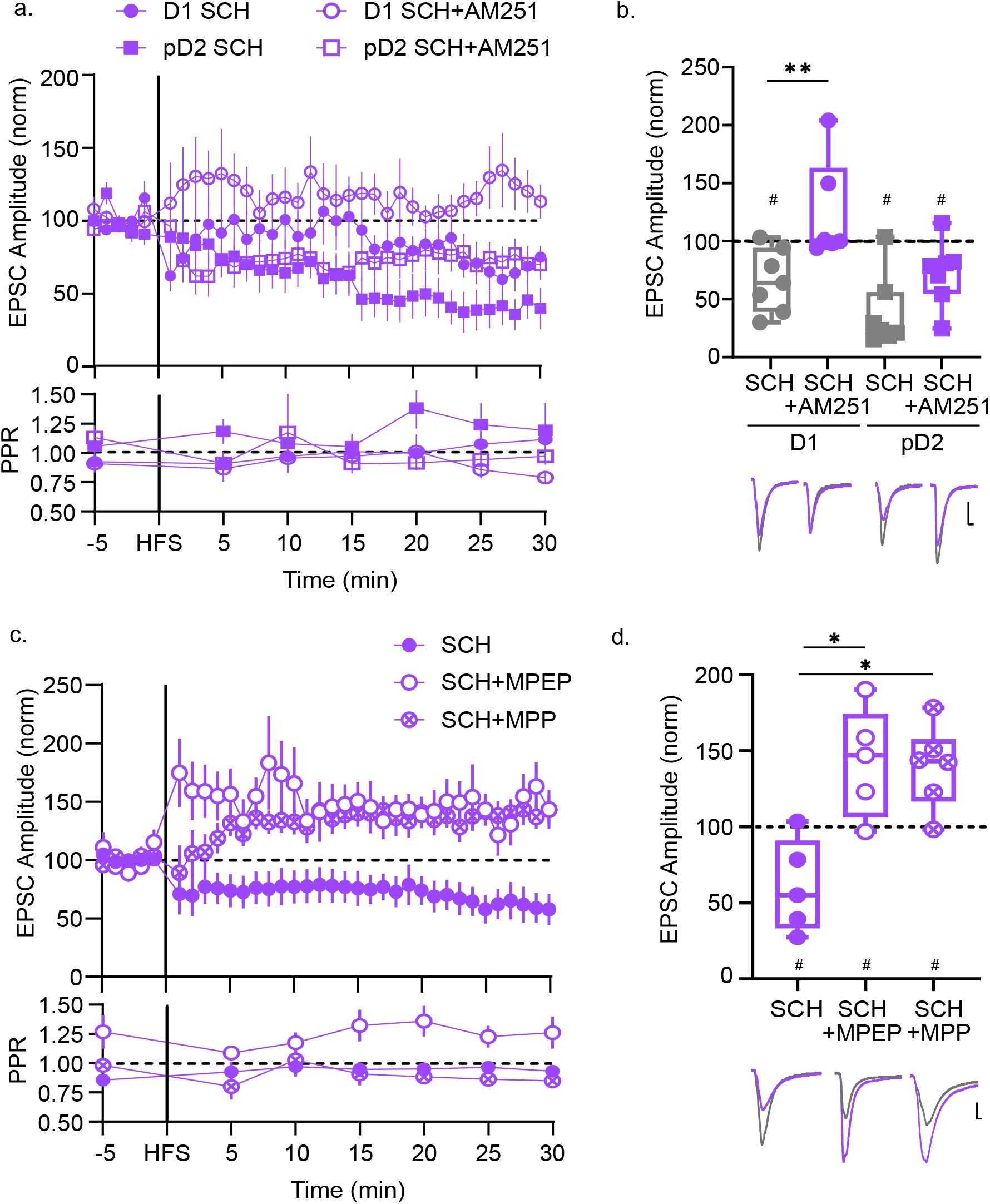
mGluR5 and ERa activity is required for SCH-LTD. a) EPSCs and PPR before and after HFS in slices pretreated with SCH50911 or SCH50911+AM251 in D1- and pD2-MSNs from females. b) Average EPSC amplitude post-HFS with representative traces. (D1 SCH n=7 cells/7 mice, W=-26, ^#^p=0.0312; D1 SCH+AM251 n = 6 cells/6 mice; pD2 SCH n=7 cells/5 mice, W=-26, ^#^p=0.0312; pD2 SCH+AM251 n=7 cells/7 mice W=-26, ^#^p=0.0312). Two-way ANOVA:F_(cell-type)_=6.668, F_(drug)_=13.88; Fisher’s LSD:*p_(D1 SCH vs SCH+AM251)_=0.0034. c) Comparison of SCH-LTD in SCH, SCH+MPEP, and SCH+MPP dihydrochloride. d) Average EPSC amplitude post-HFS with representative traces. (SCH n=5 cells/5 mice, t=0.874, ^#^p=0.0453; SCH+MPEP n=5 cells/5 mice t=2.819, ^#^p=0.0479; SCH+MPP n=6 cells/5 mice t=3.584, ^#^p=0.0158; One-way ANOVA:F=11.97 Holm-Šídák’s:*p_(SCH vs SCH+MPEP)_=0.0018, *p_(SCH vs SCH+MPP)_=0.0018). Scalebars: 20pA/10ms.

We next examined the contribution of group I mGluRs as they are typically involved in eCB-dependent LTD[16]. We found that inhibition of mGluR5 with MPEP abolished SCH-LTD at Hipp-NAc synapses (Figure 3c-d), demonstrating that mGluR5 activity is required for SCH-LTD in both MSN subtypes.

In the NAc, membrane-associated ERαs have been shown to functionally couple with mGluR5 to regulate intracellular signaling and synaptic plasticity[17-18]. Because we previously demonstrated that ERα is necessary for Hipp-NAc LTP in females[14], we asked whether ERα also contributes to the mGluR5-dependent mechanisms underlying SCH-LTD. Pretreatment of slices with ERα antagonist MPP dihydrochloride and SCH restored HFS-induced LTP (Figure 3c-d), indicating that ERα activity is required for SCH-LTD. Together, these results identify a signaling cascade in which GABA_B_R inhibition unmasks an ERα- and mGluR5-dependent form of LTD that engages presynaptic CB1R mechanisms in D1-MSNs in females.

## 4. Discussion

Our findings reveal a previously unrecognized role for GABA_B_Rs in gating the direction of synaptic plasticity at Hipp-NAc synapses in a sex-specific manner. In females, disrupting GABA_B_R signaling was sufficient to convert HFS-induced LTP into LTD without altering basal transmission, indicating that GABA_B_Rs act as key regulators of plasticity beyond their previously described role as tonic regulators of basal excitatory drive.

Mechanistically, our data indicate that this form of plasticity engages ERα and mGluR5 signaling and is expressed presynaptically at Hipp-D1-MSN synapses. Increasing evidence suggests that estradiol can rapidly influence excitatory synaptic function[19], and one major mechanism by which this has been shown to occur in the NAc is through interactions between membrane-associated ERα and mGluR5[17-18]. Taken together with our previous work demonstrating that ERα is also required for Hipp-NAc LTP in females[14], this highlights ERα as a key factor regulating plasticity of these synapses in females. This convergence between neuroendocrine and glutamatergic signaling may provide a molecular substrate through which estradiol-dependent mechanisms influence the direction of experience-dependent synaptic plasticity.

A key remaining question is where GABA_B_Rs are acting to elicit these effects. In the NAc core of male mice, GABA_B_ heteroreceptors expressed presynaptically on glutamatergic terminals are key targets for parvalbumin-expressing interneurons acting within feedforward inhibitory circuits to modulate glutamatergic transmission onto MSNs[6]. However, our wash-on experiment suggests that GABA_B_R influence on Hipp-MSN plasticity in females is likely not due to modulation of presynaptic function. Importantly, sex differences in GABA_B_R distribution and signaling have been reported in several brain regions[20-21]. Further investigation in both sexes will be necessary to determine the cellular and circuit mechanisms through which GABA_B_Rs differentially regulate plasticity in the NAc.

Together, these findings reveal a previously unrecognized, sex-specific role for GABA_B_Rs in regulating Hipp-NAc synaptic plasticity. Given previously established behavioral roles for Hipp-NAc synapses in contextual reward learning and motivated behavior[8,10,22], GABA_B_R-dependent regulation of plasticity direction may influence how Hipp information is incorporated into reward-related behaviors in females. This mechanism provides a promising avenue for further investigation into sex differences in reward circuitry, behavior, and psychiatric disorders.

## Acknowledgements

This work was supported by NSF IOS2402645, T32GM144876-02, and start-up funds from UMBC. The authors report no conflicts of interest.

